# Stable biphasic interfaces for open microfluidic platforms

**DOI:** 10.1101/392258

**Authors:** Ulri N. Lee, Jean Berthier, Jiaquan Yu, Erwin Berthier, Ashleigh B. Theberge

**Author notes:** Email address for correspondence: Ashleigh Theberge.

## Abstract

We present an open microfluidic platform that enables stable flow of an organic solvent over an aqueous solution. The device features apertures connecting a lower aqueous channel to an upper solvent compartment that is open to air, enabling easy removal of the solvent for analysis. We have previously shown that related open biphasic systems enable steroid hormone extraction from human cells in microscale culture and secondary metabolite extraction from microbial culture; here we build on our prior work by determining conditions under which the system can be used with extraction solvents of ranging polarities, a critical feature for applying this extraction platform to diverse classes of metabolites. We develop an analytical model that predicts the limits of stable aqueous-organic interfaces based on analysis of Laplace pressure. With this analytical model and experimental testing, we developed generalized design rules for creating stable open microfluidic biphasic systems with solvents of varying densities, aqueous-organic interfacial tensions, and polarities. The stable biphasic interfaces afforded by this device will enable on-chip extraction of diverse metabolite structures and novel applications in microscale biphasic chemical reactions.

---

Multiphase flows in traditional closed microfluidic channels have been harnessed for applications including extraction^1,2,3,4,5^ and interfacial chemical reactions.^6,7,8^ Our work here contributes to the broader field of open microscale methods in which fluids are manipulated in microwells, channels, or on surfaces with at least one face open to the external world.^9,10,11,12^ Open systems eliminate device-to-world leak tight connectors required by closed channel microfluidics, therefore reducing fabrication requirements. Moreover, operation of open microscale platforms is accomplished by simple pipetting, rather than sophisticated pumping systems employed by traditional closed channel microfluidics.^13,14,15^ Simple external support equipment enables increased translation and accessibility of open microfluidic devices. Here, we extend the potential of biphasic microfluidics to a simpler class of open microscale devices where passive forces are used to enable stable biphasic flows.

Manipulating biphasic flows in open microfluidic systems is challenging and requires consideration of numerous factors including air-liquid interfacial tension, aqueous-organic interfacial tension, relative wetting of the aqueous and organic phases on the device material, and Laplace pressure differences created by curved interfaces (a function of device dimensions and the volume of liquid added). Harnessing the properties of a novel class of flow we call ‘suspended microfluidics,’ where fluids flow in channels devoid of a ceiling and floor or over a floor containing apertures, we have previously developed open biphasic liquid-liquid extraction systems for isolation of steroid hormones from microfluidic human cell cultures^12^ and secondary metabolites from microscale microbial cultures.^16^ Our prior suspended microfluidic extraction systems^12,16^ were designed specifically for the properties of the solvents in those studies; we did not develop general design rules to enable the translation of the device to additional solvents. Further, the device design employed in Barkal et al. did not allow stable aqueous-organic interfaces when using solvents denser than water (e.g., chloroform).^16^ In the Barkal et al. platform, chloroform could only be employed to extract metabolites from solid agar cultures, but not from liquid cultures because the solvent was in direct contact with the culture area; in contrast, in the present manuscript the organic solvent only contacts the aqueous phase through defined microscale apertures (125-800 µm). Here, we aim to expand and generalize microscale biphase extraction systems by providing design rules for others to implement suspended microfluidic features into their devices with their choice of aqueous solution, device material, and organic solvent, including solvents denser than water. In the present manuscript, we validate an analytical model that relates the contact angle and interfacial tension of organic solvents to the stability of the biphasic interface, enabling the material and reagent properties desired for an application to drive the device design.

Liquid-liquid extraction is a widely used sample preparation technique for analysis in liquid chromatography-mass spectrometry (LC-MS)^17,18,19^ and capillary electrophoresis.^20,21,22^ The ability to work with diverse organic solvents is critical for metabolite extraction since metabolites span a range of polarities, requiring adjustment of the extraction solvent depending on the partition coefficient(s) of the analyte(s) of interest.^23,24,25^ Therefore, when designing microscale liquid-liquid extraction systems, it is essential to develop a method adaptable to solvents of varied polarities for optimal extraction efficiency.

Here, we advance our previously established suspended microfluidic biphasic extraction method by developing design rules to extend this device to solvents typically used in liquid-liquid extraction that span a range of polarities and organic-aqueous interfacial tensions. We present an analytical model of the pressure balances required to obtain stable organic-aqueous flows in devices with suspended microscale apertures connecting a lower aqueous channel and an upper organic solvent compartment. We streamline the physical factor considerations into generalized rules to enable others to easily tune their device design to work with a chosen organic solvent. Our simplified equations relate aperture radius, aqueous-organic interfacial tension, and contact angle to guide the design of open biphasic devices for optimum aqueous-organic interface stability. Conventional macroscale biphasic extractions rely on the density of the two phases to determine whether the analyte of interest will extract up into the organic phase (ρ_organic_ < ρ_aqueous_) or down into the organic phase (ρ_organic_ > ρ_aqueous_).^26^ With our design rules, we show that the effect of fluid density is negligible–enabling denser solvents such as chloroform to flow stably on top of an aqueous phase. Our model lays the foundation for biphasic open microfluidics across a range of applications including small molecule extraction and interfacial chemical reactions.

## EXPERIMENTAL SECTION

### Device Fabrication

Devices were fabricated through Midwest Prototyping (Blue Mounds, WI) using stereolithography. Devices were printed out of Accura→ 25 resin in high resolution with a z-compensation factor set to 0, level 1 finishing, and the top of the device facing up. Pressure sensitive adhesive (MicroAmp^®^ Optical Adhesive Film, Applied Biosystems^®^) was applied to the backside of the device to create the floor of the aqueous channel. Then, the devices were oxygen plasma-treated at 100 W for 50 s, with an oxygen flow rate of 5 standard cubic centimeters (sccm) (FEMTO; Diener Electronic). After plasma treatment, the tops of the devices were gently wiped with a KimWipe to prevent unwanted wetting of the tops of the device.

### Cell Culture Media

The aqueous phase was prepared by mixing cell culture media and red food coloring in a 20:1 volumetric ratio. The cell culture media (designed for NCI-H295A cells) consisted of phenol red-free DMEM/F-12 medium (Sigma–Aldrich) containing 5% (vol/vol) Nu-Serum I (BD Biosciences), 1% ITS+ Premix (BD Biosciences), 100 U/mL penicillin, and 100 μL/mL streptomycin. Red food coloring contained water, glycerine, RD&C Red #40, citric acid, and sodium benzoate.

### Organic Solvent

The organic phase was prepared by mixing Solvent Green 3 (Sigma–Aldrich) and chloroform (Fischer Chemical) to make a 18.96 mg/mL stock solution. This stock solution was diluted 20X for photos and videos in Figure 1D and diluted to 400x for Figure 1B-C. Figure 2 utilizes chloroform without dyes.

**Figure 1.**
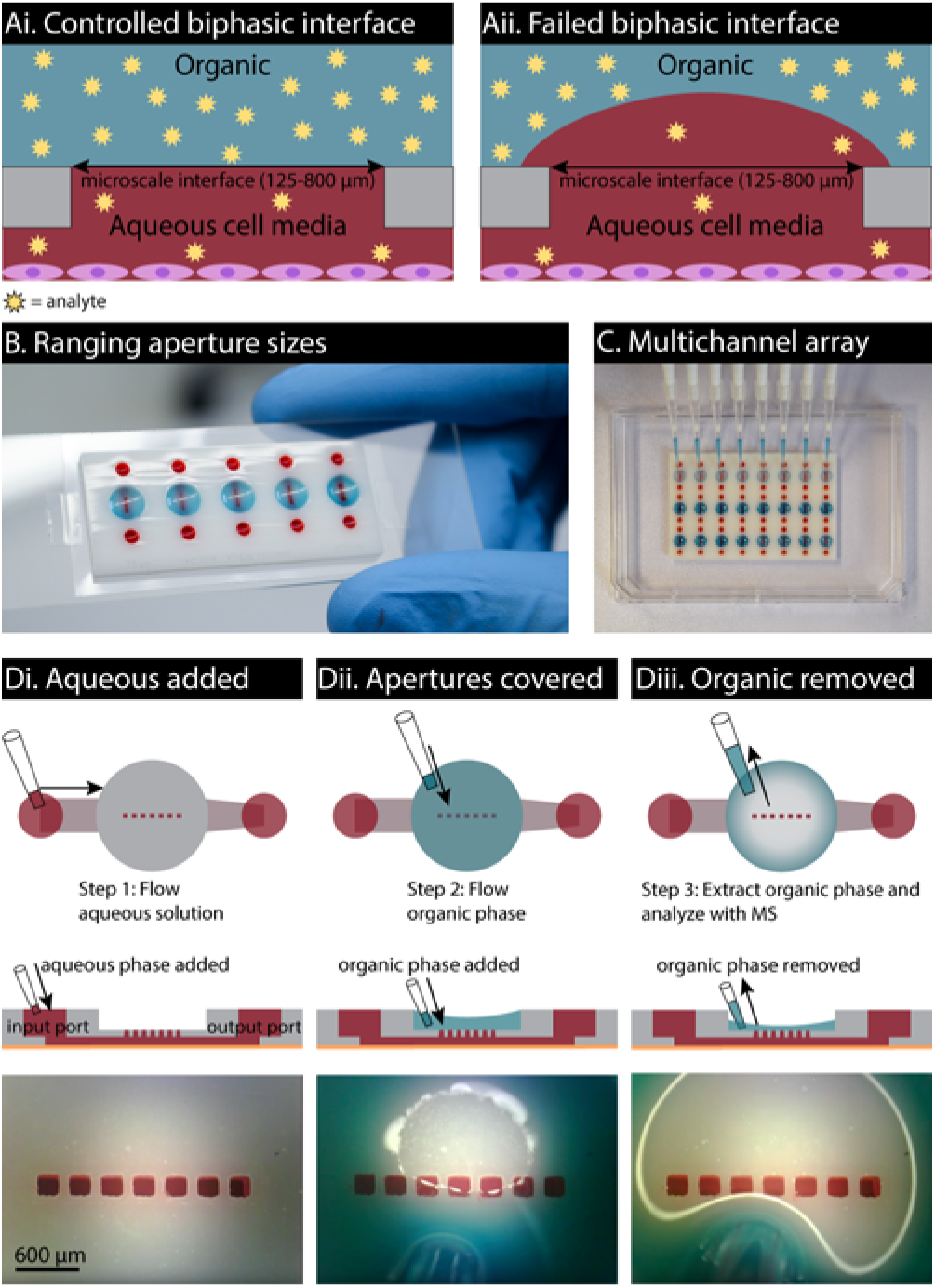
Design and operation of biphasic open microfluidic analyte extraction device. (A) The design enables addition of aqueous sample (such as cell culture, blood, etc.) in the lower channel and stable suspended flow of the organic phase over the aqueous interface. (Ai) Successful analyte extraction and (Aii) contamination of the organic phase with aqueous sample. (B) Device with port sizes ranging from 125 to 800 μm filled with cell culture media (red) and chloroform (blue). (C) Open microfluidic design enables multiplexed liquid-liquid extraction using a multichannel pipette. (D) Successful device operation; photos taken over time (excerpted from a movie, see SI) (Di) Bottom channel filled with aqueous media. (Dii) Chloroform pipetted on top of aqueous phase, successfully covering apertures. (Diii) Removal of chloroform using pipette. In this example, the red dye is not soluble in chloroform, so no small molecule extraction is observed. Device schematic provided in Figure S-1.

**Figure 2.**
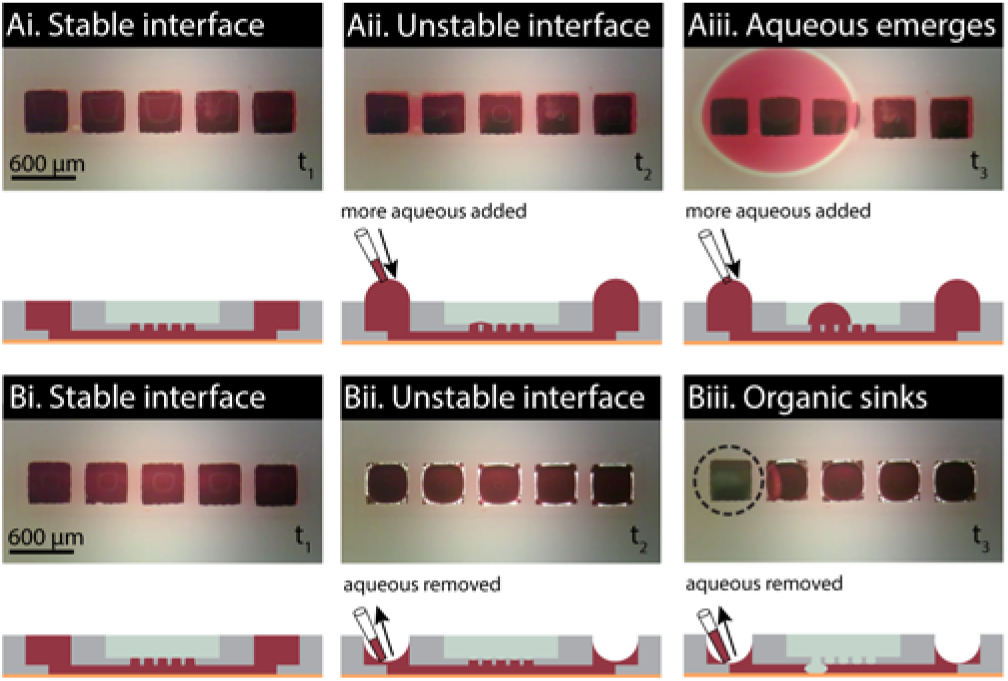
Two distinct modes of failure (destabilization of the aqueous-organic interface), resulting in contamination of the upper organic compartment with aqueous phase (cell culture media, red) (A) or the lower aqueous channel with organic solvent (chloroform, colorless) (B). Photos show top view of failed device operation over time (excerpted from a movie, see SI) with corresponding side view schematics.

### Device Operation

As shown in Figure 1D, aqueous media was pipetted with a micropipette into the input port of the aqueous channel until the aqueous-air interface in the apertures were flat. Chloroform was pipetted into the top compartment over the apertures until its organic-air interface was also flat. The chloroform was removed via pipette. The red dye is not soluble in chloroform, so no small molecule extraction was observed (the dyes used in this manuscript were selected to visualize the aqueous and organic phases separately, not to visualize extraction).

### Experimental Testing of Interfacial Stability

The following procedure was used to generate the experimental data points plotted on the phase diagrams (Fig. 4): aqueous phase was pipetted into the bottom channel such that the aqueous-air interfaces at the input/output ports were horizontal (i.e., neither concave nor convex). Solvent was pipetted into the solvent compartment, and then aqueous phase was either added (in overfilling experiments) or removed (in underfilling experiments) in 0.5-2.0 µL increments. The device was observed under a stereoscope to determine if the change in aqueous volume resulted in device failure due to instability in the aqueous-solvent interface.

**Figure 4.**
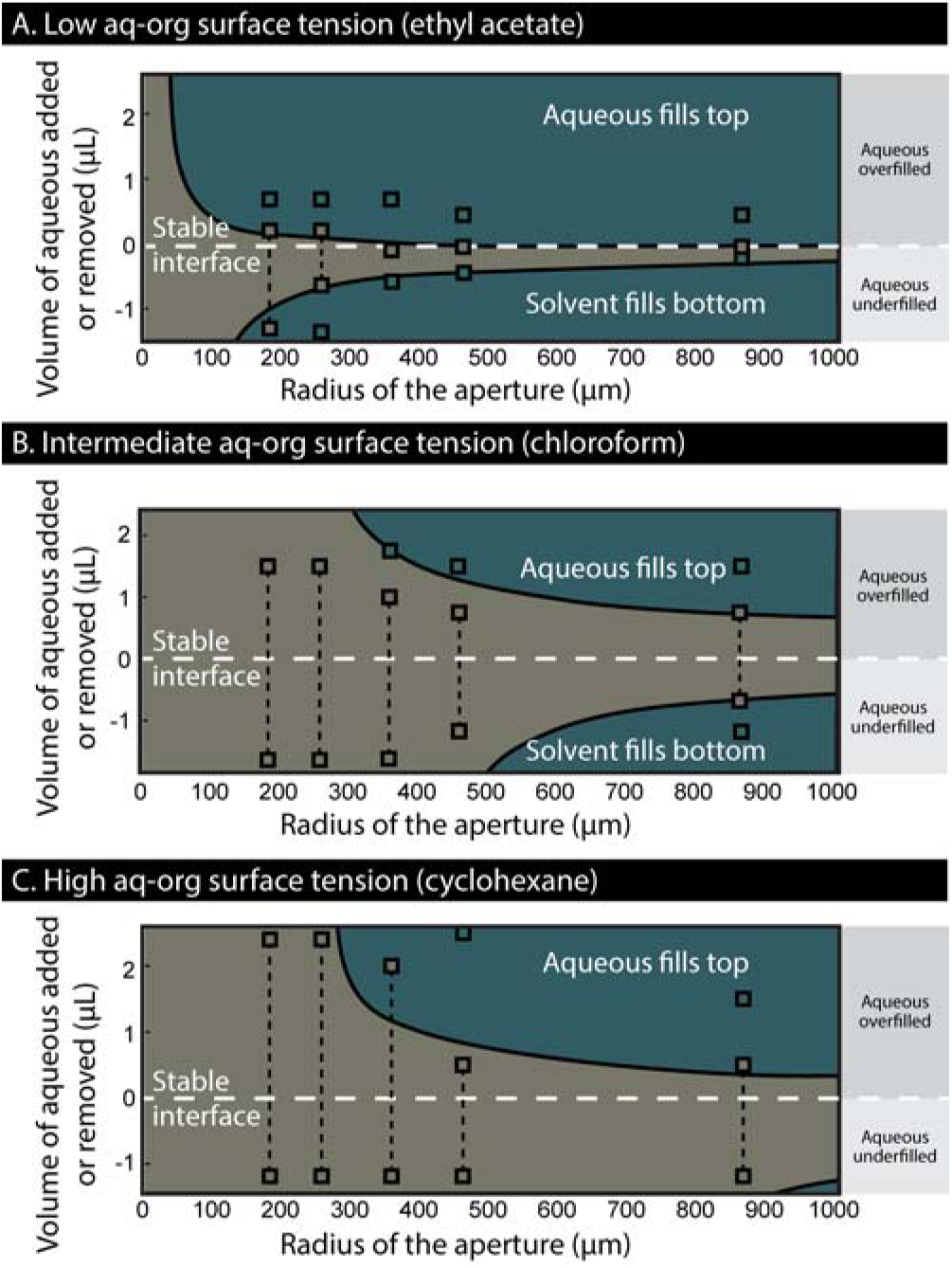
Phase diagrams showing regions of interfacial stability (gray) or failure (blue) for (A) ethyl acetate, (B) chloroform, and (C) cyclohexane across a range of measured aperture sizes (x-axis) and aqueous pressures (y-axis, indicated as aqueous volume over-or underfilled). Gray and blue shaded regions show predictions of the analytical model. Experimental results indicating interfacial stability (gray squares) or failure (blue squares) are superimposed on the predicted phase diagram, showing close correspondence between experimental results and the model. Vertical black dotted lines indicate the volumes where the interface is stable for three or more devices when aqueous solution was added or removed. Aqueous volumes were added or removed in 0.5-2.0 µL increments. In all cases, experiments were repeated with at least three microfluidic devices. Plotted points indicate volumes at which the devices were consistently stable or failed (as indicated) across all microfluidic device replicates.

### Phase Diagram Plotting

The conditions for stability were calculated based on Equation 2. We measured θ_aq,org,plastic_ experimentally with a Ramé-Hart goniometer (Netcong, New Jersey, USA) for the three solvents: approximately 40° for chloroform and 25° for ethyl acetate and cyclohexane. The parameter space was calculated as a function of volume and the radius of the apertures. The volume was used as it is directly accessible experimentally. For underfilling experiments, a pressure difference across the interface was not experienced in the system until the interface reached the bottom of the nozzle, thus, the volumes of aqueous in the nozzles were subtracted from the experimental volumes removed in the underfilling experiments. The volume thresholds corresponding to the Laplace pressure thresholds detailed in Equation 2 were calculated using trigonometric relations in a spherical cap relating the radius of curvature to the volume of the cap and the radius of the aperture. The MATLAB files used to plot the analytical model and the experimental data are included in the SI files. We limited the range of overfilling and underfilling volumes to those that ensured that the spherical cap was bound by the port of the device (when the spherical cap’s volume goes over a hemispherical cap (>90°) it will spill over the port; if it goes below an angle of −70° it will recede into the port).

## RESULTS AND DISCUSSION

We developed an open microfluidic platform that enables stable flow of an organic solvent over an aqueous sample to extract hydrophobic metabolites from the aqueous sample into the organic phase (Figure 1). The overall concept of liquid-liquid extraction across a controlled microscale biphasic interface of aqueous sample and organic solvent is shown in Figure 1Ai. Our goal here was to determine device design and operation parameters allowing stable aqueous-organic interfaces across a range of organic solvents. One potential mode of device failure we chose to investigate was the destabilization of the interface by the pressure in the aqueous phase (Figure 1Aii). We designed a device with simple geometries to accurately predict the surface-tension induced pressures throughout the channel. The device includes two stacked compartments, one below for aqueous media and a second above for organic solvent. The two compartments are connected by square apertures that stabilize the biphasic interface. The device was arrayed to test a range of aperture sizes connecting the two compartments with dimensions ranging from 125 to 800 μm (Figure 1B). The 3D printed microscale open biphasic device includes open aqueous ports and an open organic solvent compartment enabling easy access with a pipette. The channels can be arrayed for use with a multichannel pipette for fast and efficient extractions (Figure 1C). Figure 1 Di-Diii shows successful operation of a device with an aperture size of 200 μm. To model liquid-liquid extraction on our platform, we pipetted cell culture media dyed red into the input port (Figure 1Di) and chloroform dyed blue into the solvent compartment until the apertures were covered (Figure 1Dii) making sure all air-liquid interfaces were flat. Successful trials were characterized by removal of the organic phase without either phase crossing over into the other compartment (Figure 1Diii).

Ethyl acetate, chloroform, and cyclohexane were used to demonstrate the applicability of this technology because of their range of dielectric constant (an indicator of solvent polarity), density, and interfacial tension values (Table 1). Stable device operation (Figure 1D) was always observed when the aqueous-air interfaces at the input and output ports were flat for the organic phases in Table 1 using 125-800 µm apertures. However, we desired to evaluate the device when the aqueous-organic interfaces were unstable; this is important for the usability of the device since it is unrealistic to require the user to maintain a precisely flat aqueous-air interface during the operation of the device. Two modes of failure were observed when operating the device by incrementally adding or removing aqueous media (Figure 2A, aqueous media bulging up into the top compartment filled with chloroform, and Figure 2B, chloroform invading the bottom channel containing the aqueous media, videos included in SI). One mode, where the aqueous phase came up into the organic solvent compartment, was observed when excess aqueous phase was pipetted through the input port (Figure 2Aiii). The other mode of failure was observed when aqueous phase was removed, and the organic phase submerged into the aqueous phase channel (Figure 2Biii).

**Table 1.** Dielectric constant, density, and interfacial tension of solvents used in overfilling and underfilling experiments.

| Organic solvent | Dielectric constant at 20°C | Density at 25°C (g/mL) | Interfacial tension (mN/m) |
| --- | --- | --- | --- |
| Ethyl acetate | 6.080<sup>a</sup> | 0.902 | 6.8 |
| Chloroform | 4.810<sup>a</sup> | 1.492 | 32.8 |
| Cyclohexane | 2.023<sup>b</sup> | 0.779 | 50.2 |

Dielectric constant data<sup>27,28</sup> adopted from <sup>a</sup>Millipore Sigma and <sup>b</sup>Maryott *et al.* All density data<sup>27</sup> adopted from Millipore Sigma. Interfacial tension values<sup>29</sup> between solvent and water were experimentally determined by Freitas *et al.*

The main driver of instability is the pressure difference across the aqueous-solvent interface as aqueous phase is added or removed. The pressure in the aqueous phase is generated by surface tension and can be evaluated using the Young-Laplace equation (Eq. 1), where ΔP is the pressure difference between the inside and outside of a curved interface in a spherical cap, γ is the interfacial tension, and R_curvature_ the radius of curvature of the spherical interface. For simplicity, we generalized the square apertures to cylinders because the ΔP at circular interfaces is a good prediction of the ΔP approximation at square interfaces.

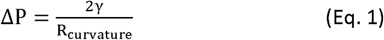

Thus, the Laplace pressure difference (ΔP) at every interface is zero when the interface is flat. When the curvature of an interface becomes concave with respect to the aqueous phase (i.e., the air-liquid interface falls below the horizontal plane), the Laplace pressure becomes negative. Inversely, when the curvature becomes convex with respect to the aqueous phase, the Laplace pressure becomes positive. The Laplace pressure is a well-defined phenomenon that can be controlled experimentally and related to the volume of liquid in the port.^15,30,31^ Thus, we were able to explore pressure ranges in the aqueous compartment (P_aq_) by varying the volume of fluid added or removed from the ports (i.e., overfilling to achieve P_aq_ > P_atm_ or underfilling to achieve P_aq_ < P_atm_). While the interface resists the pressure difference, the system is considered stable, and the two fluids remain in their respective compartments.

The pressure in the aqueous phase can eventually reach a threshold pressure (ΔP_max_) in which one of two configurations leads to the interface failing and the aqueous phase filling the solvent compartment: (1) If the angle of the aqueous-solvent interface with the horizontal line reaches 90° (i.e., the aqueous forms a hemispherical cap), the interface reaches its maximum resistance potential (maximum Laplace pressure drop) and cedes. (2) If the geometrical angle between the aqueous-solvent interface and the horizontal plane reaches the Young equilibrium contact angle between the aqueous phase, solvent, and plastic (θ_aq,org,plastic_), the aqueous media will wet the plastic and expand sideways, causing the interface to cede (Figure 3A condition (3) and Figure 2Aiii). In our experiments the contact angle θ_aq,org,plastic_ was less than 90° (θ_aq,org,plastic_ ~40°); therefore condition (2) is the limiting condition. This phenomenon of pinning at an edge, called canthotaxis, has been investigated and utilized in other systems.^4,32,33,34^

**Figure 3.**
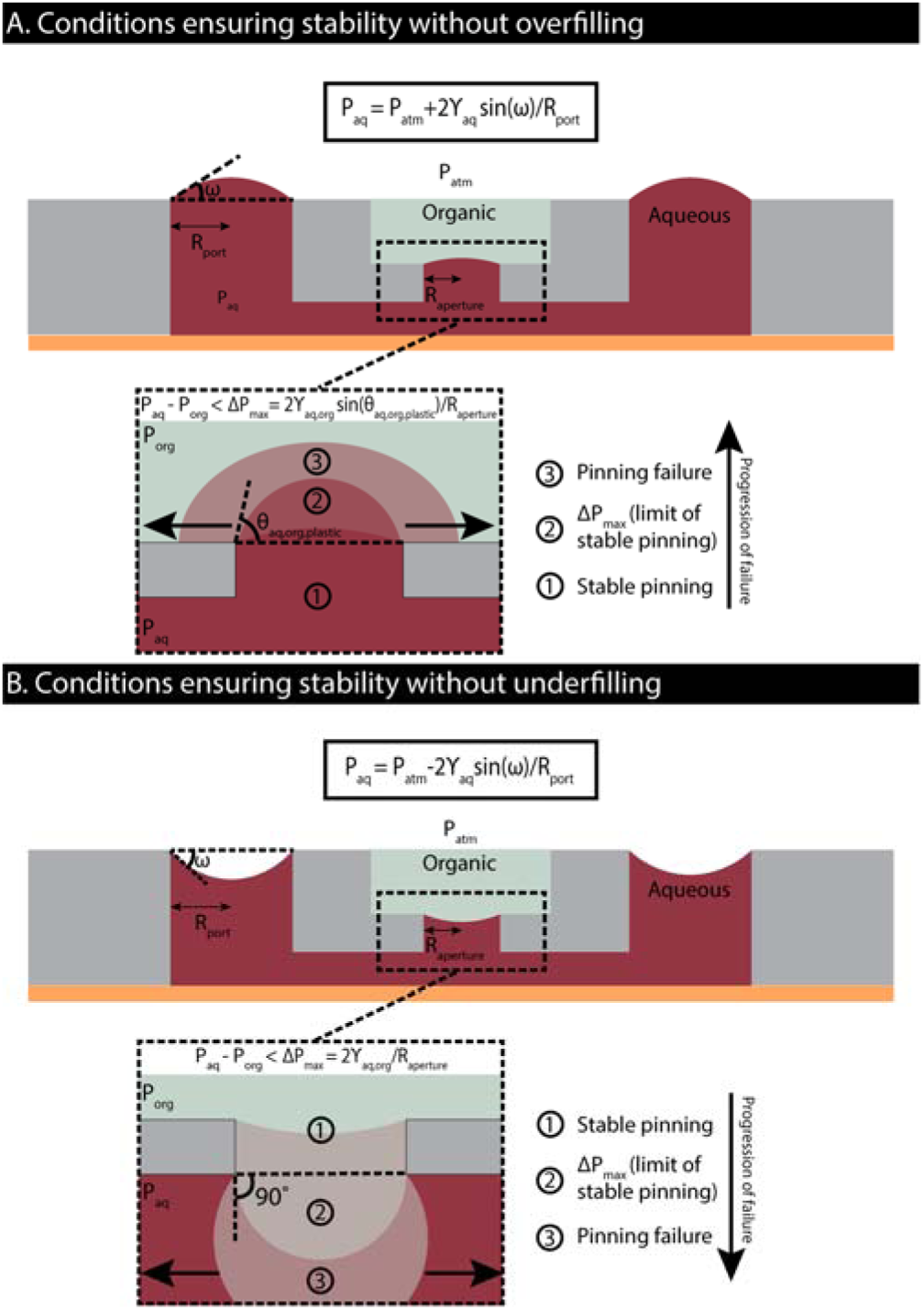
Model of Laplace pressure forces behind device failure. (A) Aqueous media is added until pinning failure, resulting in aqueous phase entering the organic phase. (B) Aqueous is removed until failure of the aqueous-organic interface, where the organic phase falls into the aqueous channel.

Using Eq. 1 and trigonometry to relate R_curvature_ to R_aperture_ (Figure S-2), we can write ΔP_max_ in terms of R_aperture_ (Eq. 2), and infer the maximum pressure difference allowed between the aqueous phase and the organic solvent to maintain stable operation of the device.

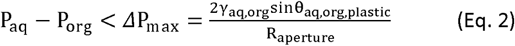

Eq. 2 is useful because it demonstrates how the aperture size can be tuned when operating the device with different solvents. Thus, as γ_aq,org_ decreases, then R_aperture_ must also decrease to allow stable device operation; this is consistent with our experimental findings (Figure 4) as described in the following section. Conversely, as aqueous phase is removed from the system, the pressure in the aqueous phase decreases. Consequently, the aqueous-solvent interface is pushed down into the aqueous compartment. The same two conditions of stability apply, by replacing θ_aq,org,plastic_ with the supplementary angle (e.g., 180°-θ_aq,org,plastic_). In our experiments θ_aq,org,plastic_ was less than 90° and the supplement was greater than 90°, thus the first condition of stability was applied (i.e., θ=90° at the limit of stable pinning) (Figure 3B). For cell culture applications where increases in temperature due to incubation would decrease surface tension, adjusted values in Eq. 2 should be used to reflect the change in the interfacial tension between the aqueous and organic phase.

We tested our analytical model by using three organic solvents of varying aqueous-organic interfacial tension: ethyl acetate (6.8 mN/m), chloroform (32.8 mN/m), and cyclohexane (50.2 mN/m).^29,35^ We plotted the boundaries of stability for varying aperture radii and the limits of over/underfilling of the aqueous media compartment for each of the solvents (Figure 4, MATLAB files available in the SI). The areas of the parameter space for which the pressure across the aqueous-solvent interface exceeded ΔP_max_ (representing device failure) are plotted in blue, and the regions of stability are plotted in gray. We validate that the devices are more stable for small aperture sizes, and the stability range decreases as the dimensions of the aperture increase. Further, the solvents with the highest interfacial tension are the most stable in this platform overall.

To test our model, we pipetted the aqueous phase into the bottom channel such that the aqueous-air interfaces at the input/output ports were horizontal by visual inspection using a stereoscope (i.e., neither concave nor convex). Solvent was pipetted into the solvent compartment, and then aqueous phase was either added (in overfilling experiments) or removed (in underfilling experiments) in 0.5-2.0 µL increments. The device was observed under a stereoscope to determine if the change in aqueous volume resulted in device failure due to instability in the aqueous-solvent interface. In Figure 4, we plotted the maximum volumes (added or removed) at which the device was stable and the minimum volumes at which the device failed. Plotted points indicate volumes at which at least three independent microfluidic devices consistently succeeded or failed (as indicated). As shown in Figure 4, the experimental results follow the analytical model and generally validate the stability conditions developed. We note that most of the discrepancy between the analytical model and experimental data occurs for small aperture sizes. This could be due to tolerance errors in 3D printing and fabrication leading to an imprecise assessment of the aperture dimensions and the generalization of square interfaces to cylinders. Further, the addition and removal of aqueous media was done by sequential pipetting events, and errors in volume can occur due to evaporation or accumulated error from multiple pipetting steps. In general, we observed a good correlation between the experimental results and the analytical predictions allowing the use of Eq. 2 as a design tool for biphasic open microfluidic device development.

## CONCLUSIONS

Our work demonstrates the ability to form controlled biphasic interfaces with water and organic solvents spanning a range of interfacial tensions and polarities, a key consideration for analyte extraction where selecting an extraction solvent with a polarity well-matched to that of the analytes of interest is critical. We have previously shown that related systems can be used to extract steroid hormones from microfluidic human cell cultures^12^ and secondary metabolites from microscale microbial cultures.^16^ Designing microfluidic devices that use passive forces to flow fluid and do not need to be connected to external pumps enables the device to move freely from incubator to bench top, which is an especially important feature when the device is both a cell culture platform and a liquid-liquid extraction device; our method adds to the growing body of self-sufficient, standalone microscale devices such as recent advances in simple reagent delivery systems.^36^ Additionally, an integrated cell culture platform streamlines the workflow required to extract molecules from cell media. Biphasic interfaces in open microfluidic devices have the potential for application outside analyte extraction such as biphasic catalytic reactions for small molecule synthesis or even utilizing biphasic reactions to create polymeric membranes^8^ or other materials at precise locations within open microfluidic channels. More fundamentally, our analytical model provides a basis for understanding the relative importance of microscale dimensions, interfacial tension, and Laplace pressure in open biphasic microfluidic devices in which different fluids are separated by pinning apertures.

## Supporting information

Fig 1D aqueous filling Fig 1D videovideo

Fig 1D organic filling video

Fig 2A overfilling video

Fig 2B underfilling video

## ASSOCIATED CONTENT

### Supporting Information

Schematic of device dimensions and mathematical relationship between θ_aq,org,plastic_, R_curvature_, and R_aperture,_ (.pdf); videos of device filling, overfilling, and underfilling from Figure 1D and Figure 2A-B (.mpg); CAD files used to 3D print devices (.sldprt) and MATLAB files for plotting conditions of stability in Figure 4 (.m) available upon request.

## ACKNOWLEDGEMENTS

This work was funded by the University of Washington, NIH K12DK100022, and NIH R01CA185251 (JY). We gratefully acknowledge Dr. David Beebe for helpful discussions and the Microtechnology Medicine and Biology (MMB) laboratory for supporting preliminary experiments that laid the foundation for this work. We thank Alexander Howard and Drs. Mark Scalf and Lloyd Smith for their contributions to preliminary experiments.

