## Supporting Information for "Stable biphasic interfaces for open microfluidic platforms"

Supporting information contents:

S-2 Schematics detailing device dimensions

S-3 Schematic of mathematical relationship between $\theta_{aq,org,plastic}$, $R_{curvature}$, and $R_{aperture.}$

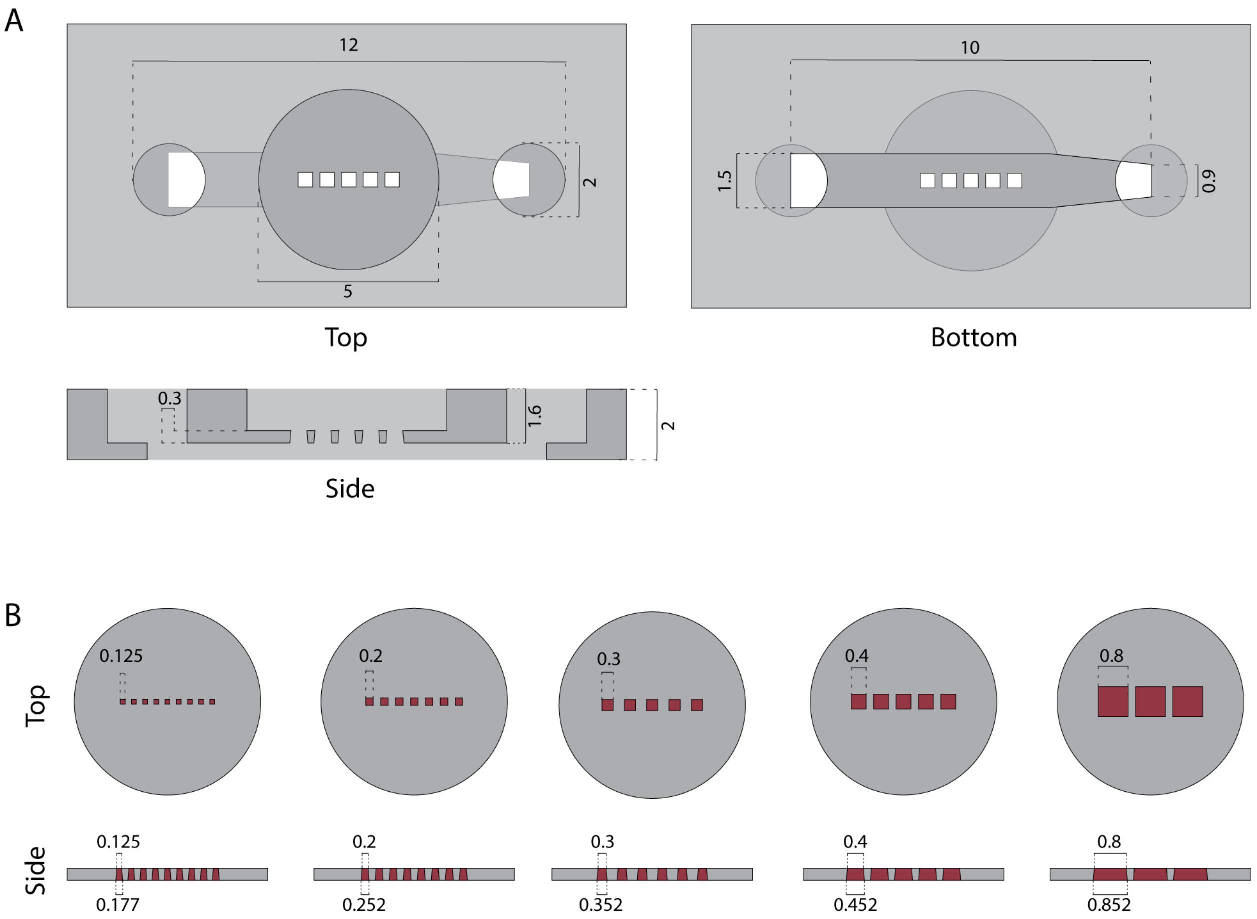


**Figure S-1**. Device schematics (A) Individual channel dimensions regardless of aperture size for open biphasic extraction device. (B) Varying aperture dimensions as seen in Figure 1B; the designed aperture dimensions are shown in this figure. Actual aperture dimensions measured were used to create the phase diagram in Figure 4 and are as follows from left to right: 0.180, 0.255, 0.355, 0.460, and 0.865 mm (apertures 3D printed larger than designed). Designed aperture sizes of 0.3 mm were used to fabricate the 8x3 array suitable for multichannel pipette use shown in Figure 1C. All measurements are shown in mm. SolidWorks files are provided as part of the SI.

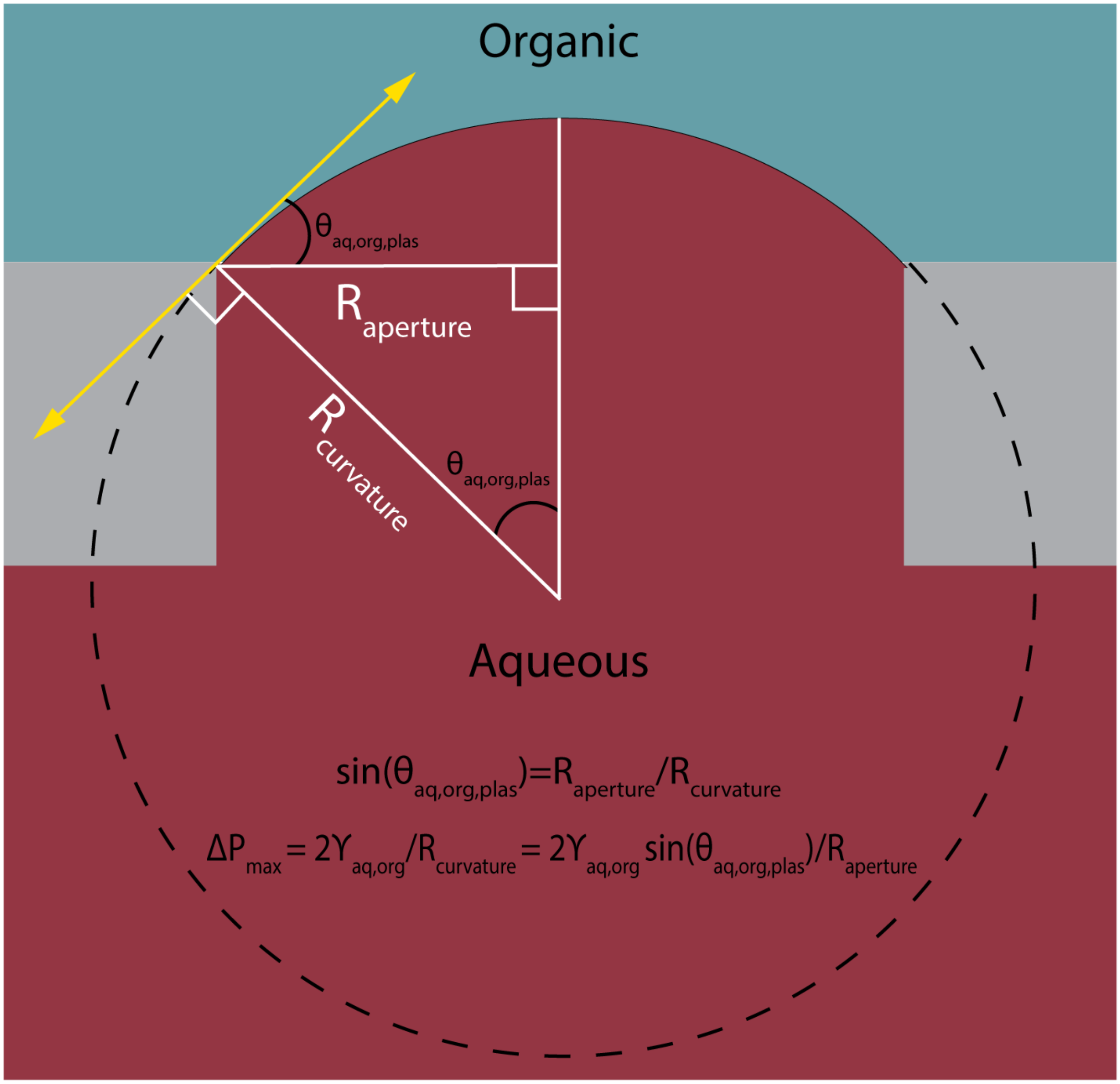


**Figure S-2.** Model of the mathematical relationship between the contact angle of the aqueous phase, organic phase, and the plastic ($\theta_{aq,org,plastic}$), the radius of curvature of the aqueous-organic interface ($R_{curvature}$), and the radius of a single aperture ($R_{aperture}$).
